# Cerebellar Gray Matter Volume Changes Across Development: Posterolateral and Vermal Transient Increases during Adolescence

**DOI:** 10.64898/2026.01.27.701939

**Authors:** Patricia Gil-Paterna, Johanna M. Hoppe, Dagmar Timmann, Richard Apps, Ebba Widegren, Matilda A. Frick, David Fällmar, Malin Gingnell, Andreas Frick

## Abstract

The cerebellum undergoes substantial maturation with regionally distinct developmental trajectories. This study examined cerebellar gray matter volume (GMV) in healthy children, adolescents, and adults, using voxel-based morphometry, the ACAPULCO algorithm, and the SUIT toolbox for cerebellum-optimized analyses. A total of 104 typically developing children (n=31, 6-9 years), adolescents (n=35, 13-17 years), and adults (n=38, 30-40 years) were included. We hypothesized age group differences in cerebellar GMV, with adolescents showing the greatest volume, specifically in posterolateral regions.

Results revealed significant group differences in GMV. We observed region-specific volumetric patterns, with some areas (e.g., Crus II, lobule X) increasing from childhood to adolescence followed by stabilization, whereas other areas (e.g., lobules I-IV and VI, Crus I, vermis VI and VIIb) exhibited peak GMV during adolescence, with lower volumes in both children and adults. These patterns were partly consistent with our hypothesis. Notably, no regions had greater GMV in adults than adolescents, suggesting that cerebellar growth peaks in adolescence before stabilizing.

Our findings indicate differential developmental patterns both between and within lobules of the cerebellum, and highlight adolescence as a peak period of cerebellar growth, with potential implications for the development of cerebellar-supported cognitive and emotional functions that undergo significant changes during this period.

## 1 Introduction

The cerebellum, nestled in the posterior cranial fossa, accounts for 10% of brain volume but contains approximately 80% of its neurons, highlighting its computational significance (Sereno et al., 2020; Azevedo et al., 2009). Traditionally linked to motor control, it is now widely recognized that its functions extend to non-motor domains, including cognition and emotion (Adamaszek et al., 2017; Middleton & Strick, 1994), via tightly organized, topographically structured architecture (Apps & Hawkes, 2009). Reciprocal connections with the neocortex, through cerebellar nuclei and thalamic relays, facilitate complex cortico-cerebellar communication supporting cognition and emotion (Schutter, 2020; D’Angelo, 2018; Bostan et al., 2013; Rudolph et al., 2023). Anatomically, the cerebellar vermis is a midline structure, lobules I-IV and V make up the anterior cerebellum and lobules VI to X are considered posterolateral regions (Schmahmann et al., 2000). Lobule VIIa is often divided into Crus I and Crus II, thus also part of the posterolateral cerebellum. For reference, Supplementary Figure 1 provides a detailed anatomical view of the cerebellar cortex, while Figure 2 (see below) illustrates an overall organization.

**Figure 1.**
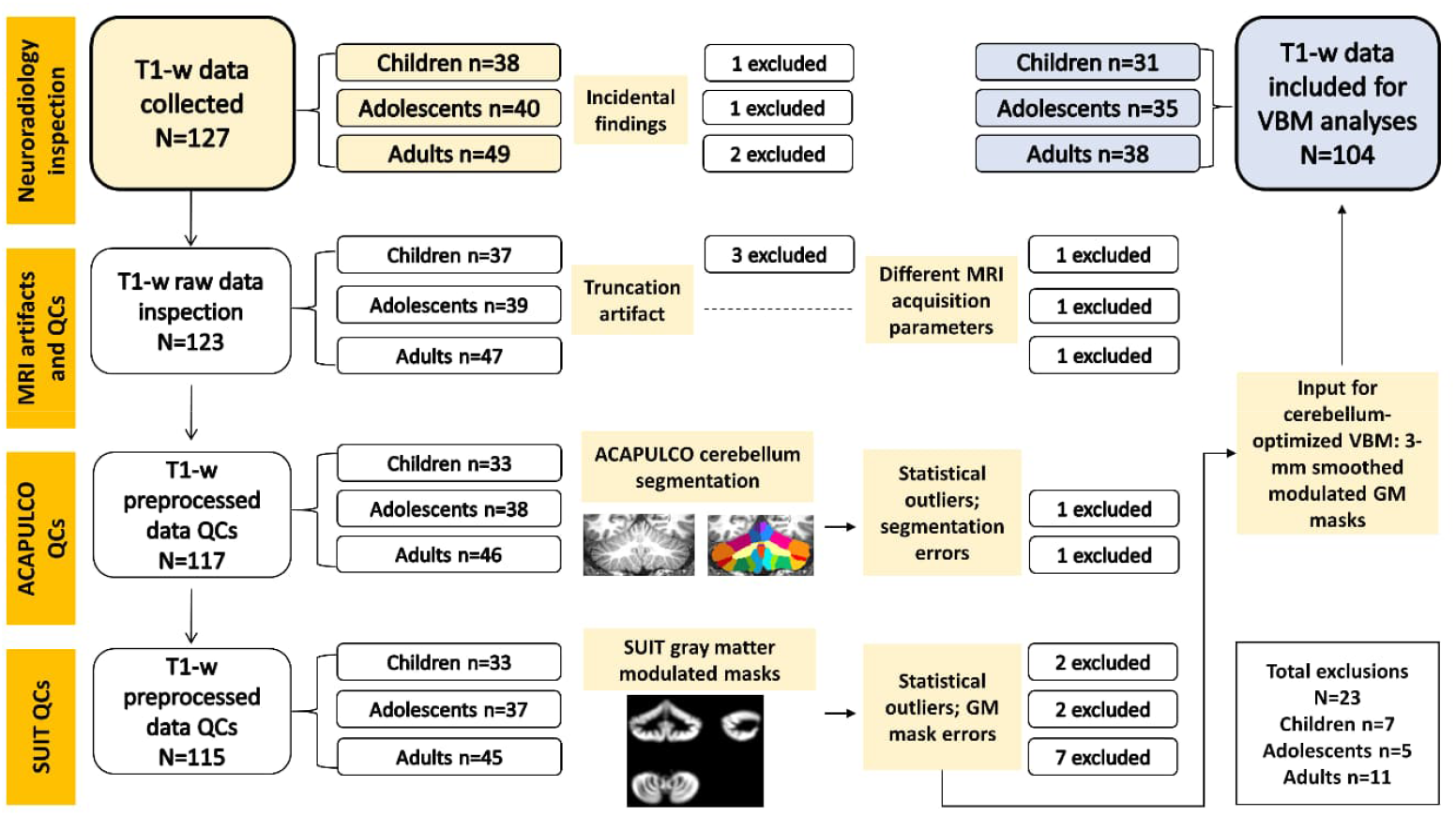
Flowchart of the exclusion criteria and quality checks (QCs) applied sequentially along the study. Out of 127 participants included in the original study, 23 were excluded, leaving 104 in the analyses. GM = gray matter.

**Figure 2.**
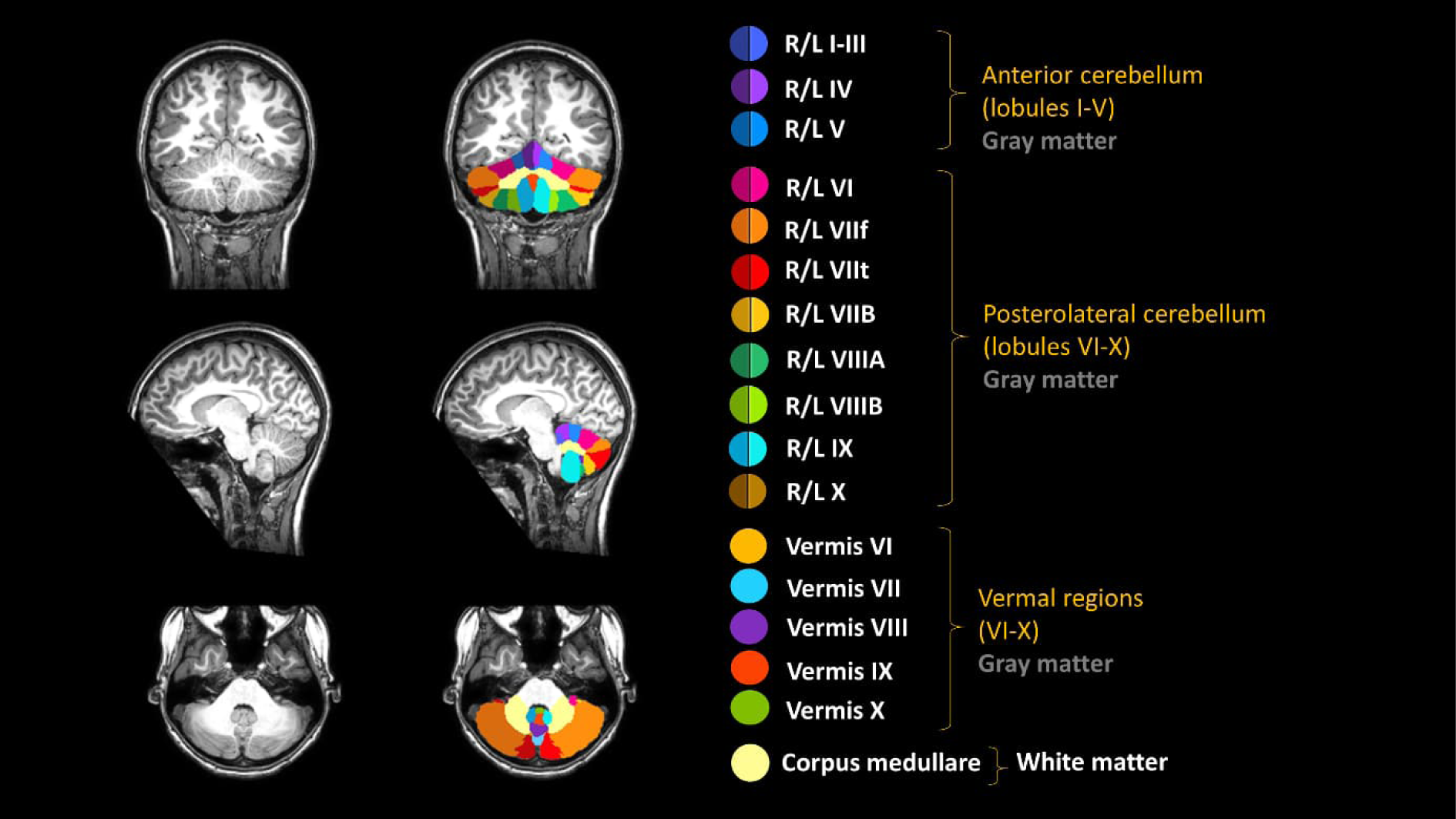
ACAPULCO cerebellum segmentation of an included adolescent. From top to bottom: coronal, sagittal, and axial views. R/L = right/left. .

The “cerebellar cognitive affective syndrome”, first described by Schmahmann and Sherman (1998), provided compelling clinical evidence for the role of the cerebellum in higher-order functions and emotion, linking cerebellar lesions to impairments in executive functioning, affect regulation, and personality. Neuroimaging studies further support this view, showing that the cerebellar vermis contribute to affective processing (e.g., Xiao et al., 2025), while more posterolateral (e.g., lobules VI, Crus I and II) and anterior regions (lobules I-V) support cognitive-emotional processes and sensorimotor functions, respectively (Guell et al., 2018a; Stoodley & Limperopoulos, 2016; Stoodley & Schmahmann, 2009). For instance, Moberget et al. (2019) found negative associations between posterior cerebellar volumes, particularly lobule VI and Crus I, and subclinical anxiety and cognitive dysfunction in healthy adolescents. More broadly, while posterolateral cerebellar regions may support cognitive-emotional integration, anterior regions and lobule VIIIb may subserve motor components of emotional processing, such as motor imagery (Habas et al., 2009). These regional specializations suggest that the cerebellum contributes to various cognitive and emotional functions, likely mediated by reciprocal loops with prefrontal regions (Ramnani, 2006; Middleton & Strick, 2001; Schmahmann & Pandya, 1997).

### 1.1 Cerebellar Morphology Across Development

Accumulating, yet limited, evidence from structural neuroimaging studies indicates that the cerebellum undergoes substantial changes across development. In an initial study, Tiemeier et al. (2010) reported that cerebellar gray matter volume (GMV) follows an inverted U-shaped trajectory from childhood to early adulthood, peaking in mid-adolescence, with region-specific timing across subregions: the inferior posterior lobe (Crus II–lobule X) peaks first, followed by the anterior lobe (lobules I–V), and finally the superior posterior lobe (lobule VI–Crus I). Subsequent studies of total cerebellar GMV yielded mixed results, with some confirming a mid-adolescent peak (Wierenga et al., 2014) and others suggesting a general age-related decline (Østby et al., 2009). More recent work using larger samples and higher magnetic fields supports that the cerebellum undergoes a non-uniform anterior-to-posterior growth gradient from childhood to adolescence (Gaiser et al., 2024; Liu et al., 2022). This growth pattern mirrors the earlier maturation of sensorimotor (localized to anterior cerebellum) compared to cognitive functions (posterior cerebellum). However, most studies have focused on total volumes within predefined lobular subdivisions or limited age ranges, leaving open questions about how cerebellar GMV evolves from childhood to mid-adulthood. Furthermore, it remains unclear whether the vermis, consistently linked to emotional behaviour, undergoes developmental changes, as prior work reported no difference in GMV between children, adolescents, and young adults (Tiemeier et al., 2010) in this small but functionally salient region. Investigating regionally specific volumetric patterns in the cerebellum is therefore crucial, as distinct subregions may differently support cognitive and affective domains (Guell et al., 2018b), and potentially contribute to the onset of developmental changes across life stages, with particular relevance during adolescence.

In this study, we examined regional cerebellar GMV in healthy children, adolescents, and adults using voxel-based morphometry (VBM) with a fully automated, cerebellum-optimized pipeline, enabling a more precise anatomical characterization of its protracted maturation. We hypothesized that GMV would significantly differ across age groups, with specific subregions showing distinct developmental patterns. Based on prior evidence (e.g., Tiemeier et al., 2010), we predicted GMV increase from childhood, peaking in adolescents, and then declining into adulthood in posterolateral regions, including lobule VI, Crus I, Crus II. In a secondary, exploratory analysis, we also investigated whether these cerebellar developmental patterns are associated with cognitive and emotional functions. By applying VBM across a wide age range from 6 to 40 years, we provide a region-specific and comprehensive perspective on cerebellar structural development, extending beyond the narrower age windows and broader anatomical specificity typically examined in prior work.

## 2. Methods

### 2.1 Participants and Procedure

Healthy, typically developing children (6-9 years), adolescents (13-17 years) and adults (30-40 years) were recruited from previous studies at the Department of Psychology, Uppsala University, Sweden, using public advertisements. The recruitment procedure has been described in detail elsewhere (Widegren et al., 2025). Briefly, potential participants were screened for eligibility through online questionnaires and a subsequent psychiatric diagnostic interview (Mini-International Neuropsychiatric Interview, Sheehan et al., 1998). Exclusion criteria included atypical development or premature birth (before 37 weeks of gestation), contraindications to magnetic resonance imaging (MRI), uncorrected vision or hearing impairment, illicit drug use or use of psychotropic medication, pregnancy, neurodevelopmental or neurological disorders, and presence or history of psychiatric disorder. A total of 127 included participants completed additional questionnaires online and underwent a two-day procedure comprising cognitive and emotional psychological tests on day 1, and neuroimaging on day 2. The study adhered to the Declaration of Helsinki and was approved by the Swedish Ethical Review Authority (2019-01929, 2022-02234-02). Written informed consent was obtained from adult participants and caregivers of minors. Children and adolescents provided informed assent.

### 2.2 Psychological Measures: Assessment of Cognition and Emotion

Cognitive performance was assessed using the Matrix Reasoning subtest of the Wechsler Intelligence Scales (Wechsler, 2014; 2008). For children and adolescents, scores were derived from the Wechsler Intelligence Scale for Children Fifth Edition (WISC–V; Wechsler, 2014). Although scores on Matrix Reasoning were available for adults, these were not included in the current study due to difference in task version (i.e., the Wechsler Adult Intelligence Scale Fourth Edition, WAIS–IV; Wechsler, 2008), precluding direct comparability. Emotional functioning was examined with the Emotion Questionnaire (EQ; Rydell et al., 2007), which measures emotional regulation and reactivity. In the present study, self-reported EQ scores from adolescents and adults only were included as the self-report version of the EQ was not administered to children.

### 2.3 MRI Acquisition

High-resolution anatomical T1-weighted (T1-w) images were acquired using a 3.0 Tesla Philips Achieva dStream whole-body MR scanner (Philips Medical Systems, Best, The Netherlands) with a 32-channel head-coil, at Uppsala University Hospital, Sweden. Acquisition parameters were as follows: repetition time/echo time/inversion time (TR/TE/TI) = 8.2 ms/3.8 ms/685.5 ms; field of view (FoV) = 240 x 240 mm, flip angle = 8°, yielding 220 contiguous slices, with voxel size = 0.94 x 0.94 x 1 mm. These parameters enabled whole-brain coverage, ensuring entire inclusion of the cerebellum. A senior consultant in neuroradiology (DF) screened all T1-w images for morphological abnormalities and/or incidental findings with possible interference with the analysis, resulting in four subjects being excluded. All images were checked for artifacts, and truncation led to three additional exclusions, whereas another three subjects were removed due to deviations in MRI acquisition parameters. A flowchart illustrating MRI quality checks and exclusion criteria is presented in Figure 1.

### 2.4 MRI Preprocessing

To enable VBM analysis, T1-w images were preprocessed using a cerebellum-optimized pipeline developed by the ENIGMA-Ataxia Working Group (Kerestes et al., 2022), the ENIGMA Cerebellum Volumetrics Pipeline (https://enigma.ini.usc.edu/ongoing/enigma-ataxia/), with custom Linux-based scripts. This pipeline uses an automatic parcellation using U-Net with locally constrained optimization (ACAPULCO) algorithm (Han et al., 2020) to locate and parcellate the cerebellum into 28 anatomical subregions, in a two-step method based on three-dimensional convolutional neural networks. Preprocessing was performed in FreeSurfer 7.4.1 (http://surfer.nmr.mgh.harvard.edu/), and included resampling T1-w images to 1x1x1 mm, and calculating estimated total intracranial volume (eTIV). Figure 2 shows an example of ACAPULCO cerebellum subsegmentation in 28 regions from an included adolescent.

### 2.5 Volumetric Analysis

Volumetric analyses were performed using the ENIGMA Cerebellum Volumetrics pipeline, with the SUIT toolbox (v3.4; Diedrichsen, 2006) in SPM12 (MATLAB R2021b, MathWorks Inc., Natick, MA). SUIT enables a cerebellum-refined VBM, in a two-step processing: isolation and tissue segmentation (step 1), and DARTEL normalization to the SUIT cerebellar template (step 2). Resulting gray matter maps were bias-corrected, MNI152-registered, and modulated by the Jacobian determinant. Quality control included checks on spatial covariance and quartic mean *z*-scores. Subjects with spatial covariance ≥ 2 SD below the mean, or with misaligned gray matter cerebellar masks were excluded. Maps were smoothed with a 3 mm Gaussian kernel, based on the pipeline description and the consideration that smaller kernels enhance the preservation of anatomical details. All participants underwent identical pre and postprocessing steps, with no manual edits to the structural MRI data.

### 2.6 Statistical Analysis

Statistical tests to assess normality, variance homogeneity, and descriptive statistics for brain volumetrics (eTIV, total cerebellum volume) as well as for scores from the psychological measures were conducted in JASP 0.19.1.0 (JASP Team, 2024), alongside ANCOVA and *t*-test analyses. Age group differences in GMV were tested using smoothed (3 mm), modulated cerebellum gray matter T1-w images, with a general linear model in SPM12. A full factorial design included eTIV (mean-centered) and sex (uncentered) as covariates. eTIV, a measure in mm^3^ of the volumetric space contained within the skull, was used to correct for different brain sizes, as commonly done in VBM analysis. To minimize potential edge effects between gray and white matter tissues, all voxels with gray matter values below 0.10 were excluded using absolute threshold masking, and explicit SUIT masking was applied. Six pairwise age group comparisons were performed. Voxel-wise results were thresholded at *p* < 0.05 Family-Wise Error (FWE)-corrected with a 20-voxel cluster extent, for greater stringency to reduce Type I error. Significant thresholded *T*-maps were visualized using SPM12 as well as FSLeyes, overlaid on the SUIT cerebellar mask. In addition to reporting MNI coordinates and *t*-values derived from VBM, Cohen’s *d* effect sizes were estimated as follows, following best-practice recommendations in neuroimaging reporting (Chen et al., 2017):

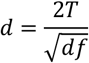

where the *t*-value multiplied by 2 (*2T*) is divided by the square root of the degrees of freedom (*df*) for each pairwise contrast [(N _age group 1_ + N _age group 2_) – 2)]. Alternatively, we calculated the eta-squared (*η*^2^) effect size measure, computing the following formula:

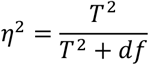

in which the squared *t*-value (*T* ^2^) is divided by the sum of the squared *t*-value and the *df* (*T* ^2^ + *df*). The effect size estimates using eta-squared provide an intuitive measure of the proportion of variance in GMV explained by age group differences at a given voxel.

Exploratory analyses were conducted in SPM12 to examine associations between cerebellar GMV and cognitive or emotional functioning. Cognitive performance (indexed by WISC-V Matrix Reasoning scores) was analyzed in relation to GMV within and between children and adolescents (adults were excluded due to completing a different version of the task). Similarly, EQ emotion regulation and reactivity scores were analyzed in adolescents and adults (children were not included as they did not complete the questionnaire). Task scores were also compared between children and adolescents (Matrix Reasoning) and adolescents and adults (EQ) using *t*-tests. All analyses included sex and eTIV as covariates and were masked using the SUIT mask. Visualization of results were achieved through regression plots from peak coordinates as well as SUIT flatmaps (Diedrichsen & Zotow, 2015). We also conducted age-invariant analyses to test for associations between cerebellum GMV and cognitive and emotional function using age, sex, and eTIV as covariates.

## 3. Results

### 3.1 Participants

Out of the 127 participants completing the data collection, 23 participants were excluded due to incidental findings (n=4), image artifacts (n=3), and acquisition (n=3) and segmentation quality issues (n=13). This left 104 participants included in the current study: 31 children (age M ± SD, 8.0 ± 0.93 years), 35 adolescents (13.7 ± 1.16 years), and 38 adults (34.4 ± 3.11 years). Figure 1 displays a visual flowchart of participant inclusion and exclusion. The ratio of female/male participants (self-reported biological sex) was 19/12 in children (61% females), 19/16 in adolescents (54% females), and 19/19 in adults (50% females). The age groups did not significantly differ in sex distribution (*χ*^*2*^= .88, *p* = .64). Total eTIV-corrected cerebellar volume was larger in adolescents than in children, with no difference detected between adolescents and adults. See the Supplementary Material for age group eTIV and total cerebellum volumes and comparisons.

### 3.2 Voxel-Based Morphometry

A general pattern of results emerged in, eTIV-corrected, voxel-wise analyses indicating an inverted U-shaped trajectory of cerebellar GMV from childhood to adulthood with a peak in adolescence (see Supplementary Figure 3). There were no clusters in which children had greater GMV than adolescents or adults, or where adults had greater GMV than adolescents.

#### 3.2.1 Adolescents > Children

Adolescents had greater GMV than children in a large cluster with its peak located in the bilateral Crus II and the cluster extending superior into Crus I, and inferior to lobules VIIb, VIIIa, VIIIb, and IX, including the vermes. More circumscribed differences were detected in bilateral lobule X, right lobule I-IV, and right Crus I. Effect sizes of these age-group differences were medium to very large. See Figure 3a and Table 1.

**Figure 3.**
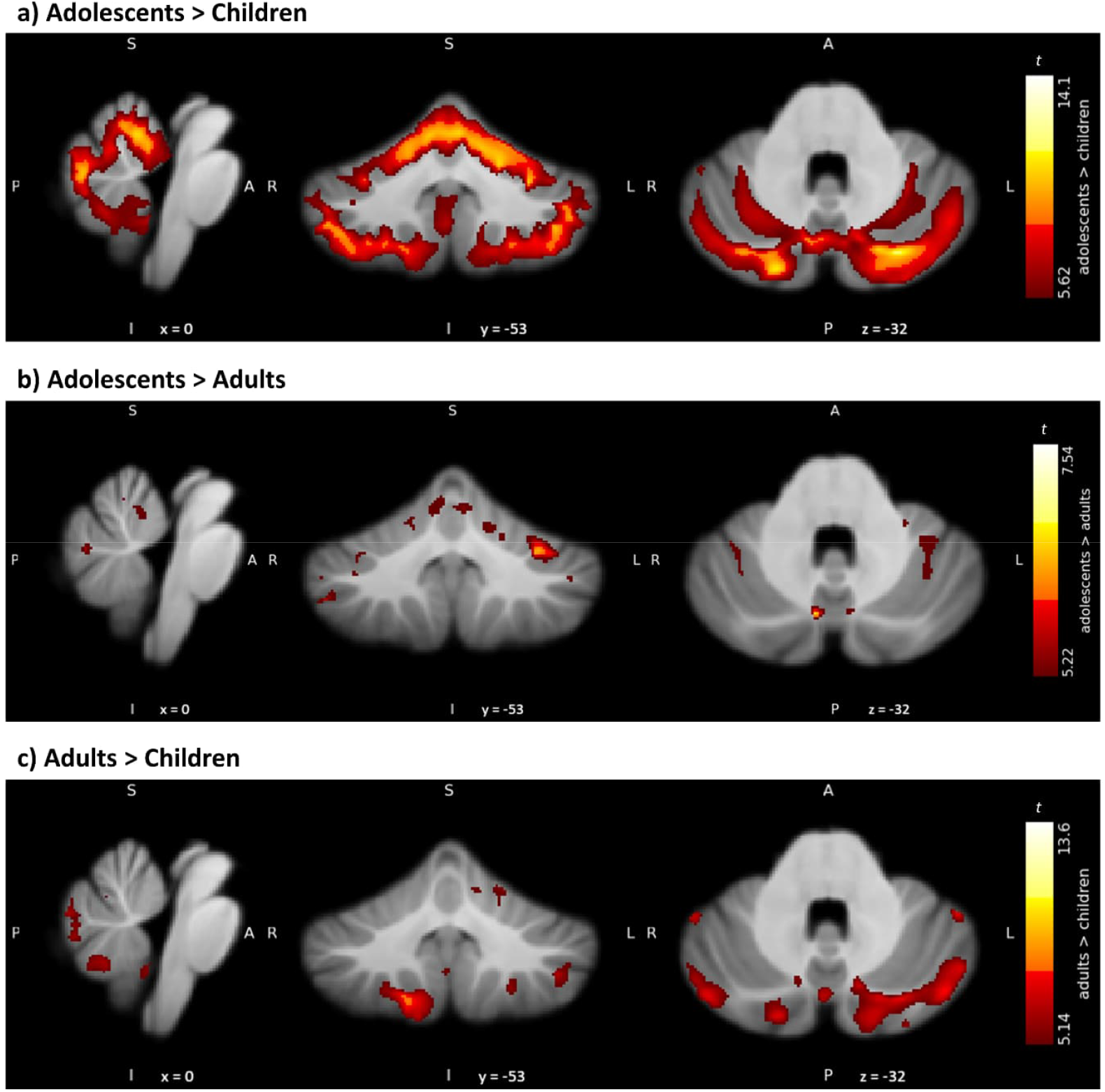
Cerebellum gray matter volume differences across age groups derived from voxel-based morphometry. **3a**) Contrast showing adolescents over children. **3b**) Contrast showing adolescents over adults. **3c**) Contrast showing adults over children. All clusters are family-wise error corrected at *P*_FWE_<0.05 and cluster extent of 20 voxels. Colorbar indicates gradient of *t*-values (*t*) in each contrast. From left to right: sagittal (x), coronal (y), and axial (z) views, with coordinates. *T*-maps were overlaid on the SUIT cerebellum template and displayed in radiological orientation.

**Table 1.** Voxel-based morphometry results comparing age groups. Thresholds set to family-wise error (FWE) corrected p<0.05 and cluster extent of 20 voxels.

| Cerebellar region | MNI coordinates | Volume (mm<sup>3</sup>) | *t* | *p*<sub>FWE</sub> | Effect size (Cohen's *d*) | $\eta^2$ (% of variance explained) |
| --- | --- | --- | --- | --- | --- | --- |
|  | x y z |  |  |  |  |  |
| Adolescents > Children |  |  |  |  |  |  |
| Left Crus II | -21 -81 -42 | 87342 | 14.01 | <.001 | 3.50 | .754 (75.4%) |
| Right X | 24 -38 -41 | 273 | 6.98 | <.001 | 1.74 | .432 (43.2%) |
| Right I-IV | 11 -43 -29 | 21 | 5.91 | .001 | 1.48 | .353 (35.3%) |
| Left X | -20 -41 -43 | 160 | 5.85 | .001 | 1.46 | .348 (34.8%) |
| Right Crus I | 48 -47 -31 | 25 | 5.62 | .002 | 1.40 | .330 (33.0%) |
| **Adolescents > Adults** |  |  |  |  |  |  |
| Vermis VI | 4 -71 -28 | 315 | 7.54 | <.001 | 1.79 | .444 (44.5%) |
| Left VI | -33 -45 -39 | 166 | 6.85 | <.001 | 1.63 | .393 (39.4%) |
| Left VI | -30 -54 -28 | 1771 | 6.36 | <.001 | 1.52 | .360 (36.0%) |
| Right Crus I | 48 -51 -46 | 136 | 6.31 | <.001 | 1.51 | .356 (35.6%) |
| Right Crus I | 49 -47 -38 | 66 | 6.08 | <.001 | 1.45 | .340 (34.0%) |
| Right Crus I | 35 -62 -36 | 255 | 5.80 | .001 | 1.39 | .316 (31.6%) |
| Right I-IV | 6 -51 -14 | 1157 | 5.74 | .002 | 1.37 | .311 (31.2%) |
| Right Crus I | 39 -45 -39 | 57 | 5.51 | .004 | 1.32 | .299 (29.9%) |
| Left VIIb | -12 -63 -44 | 26 | 5.22 | .011 | 1.25 | .276 (27.6%) |
| **Adults > Children** |  |  |  |  |  |  |
| Left Crus II | -21 -81 -42 | 22893 | 13.63 | <.001 | 3.32 | .735 (73.5%) |
| Right Crus II | 27 -80 -43 | 13807 | 12.07 | <.001 | 2.93 | .684 (68.4%) |
| Right I-IV | 6 -38 -17 | 284 | 7.89 | <.001 | 1.91 | .481 (48.1%) |
| Left IX | -11 -43 -44 | 1139 | 7.70 | <.001 | 1.89 | .471 (47.1%) |
| Left Crus I | -47 -45 -33 | 67 | 7.43 | <.001 | 1.82 | .454 (45.4%) |
| Right VI | 20 -72 -19 | 786 | 7.17 | <.001 | 1.76 | .432 (43.2%) |
| Right IX | 12 -42 -48 | 144 | 6.91 | <.001 | 1.68 | .418 (41.8%) |
| Right Crus I | 51 -47 -32 | 61 | 6.69 | <.001 | 1.63 | .396 (39.7%) |
| Left X | -20 -41 -44 | 183 | 5.93 | .001 | 1.44 | .341 (34.1%) |
| Right V | 1 -63 -18 | 29 | 5.43 | .005 | 1.32 | .303 (30.3%) |
| Right I-IV | 11 -48 -16 | 36 | 5.39 | .006 | 1.31 | .298 (29.9%) |
| Right V | 7 -61 -22 | 199 | 5.27 | .009 | 1.28 | .286 (28.6%) |
| Right I-IV | 1 -47 -18 | 24 | 5.14 | .015 | 1.25 | .277 (27.8%) |

MNI = Montreal Neurological Institute.

#### 3.2.2 Adolescents > Adults

Adolescents exhibited greater GMV than adults in a number of widespread clusters with medium to large effect sizes, including bilateral lobules I-IV and VI, extending to the inferior posterolateral regions such as Crus I and left VIIb, whereas vermis VI and VIIb formed a distinct cluster. See Figure 3b and Table 1.

#### 3.2.3 Adults > Children

Adults had greater GMV than children, mostly in the anterior and posterolateral regions of the cerebellum with the two largest clusters extending from bilateral Crus II. Effect size estimates were large to very large. See Figure 3c and Table 1.

### 3.3 Cerebellar Contributions to Cognitive and Emotional Function

For the WISC-V Matrix Reasoning subtest, children scored lower (16.48 ± 3.29 raw points) than adolescents (22.46 ± 2.94 raw points), Student’s *t*(64) = -7.79, *p* < .001, Cohen’s *d* = -1.92. Scores were normally distributed and variances were homogeneous (Saphiro-Wilk *W* = 0.96, *p* = .09; Levene’s *F* = 1.69, *p* = .19).

On the EQ Emotional Regulation scale, adolescents rated lower emotion regulation capacity (3.48 ± 0.79) than adults (3.83 ± 0.58), Welch’s *t*(59.93) = -2.14, *p* = .036, *d* = .51. Although the data deviated from normality (*W* = 0.94, *p* = .003), variances were equal (Levene’s *F* = 0.60, *p* = .44). We detected no difference in emotional reactivity between adolescents (2.27 ± 0.76) and adults (2.48 ± 0.67), Welch’s *t*(66.10) = -1.22, *p* = .23, *d* = -0.29. Data were slightly non-normal (*W* = 0.97, *p* = .046; Levene’s *F* = 0.80, *p* = .37). Q-Q plots of standardized residuals were checked for data being not normally distributed, indicating only minor outliers. Figure 7 displays raincloud plots of cognitive and emotional scores.

**Figure 7.**
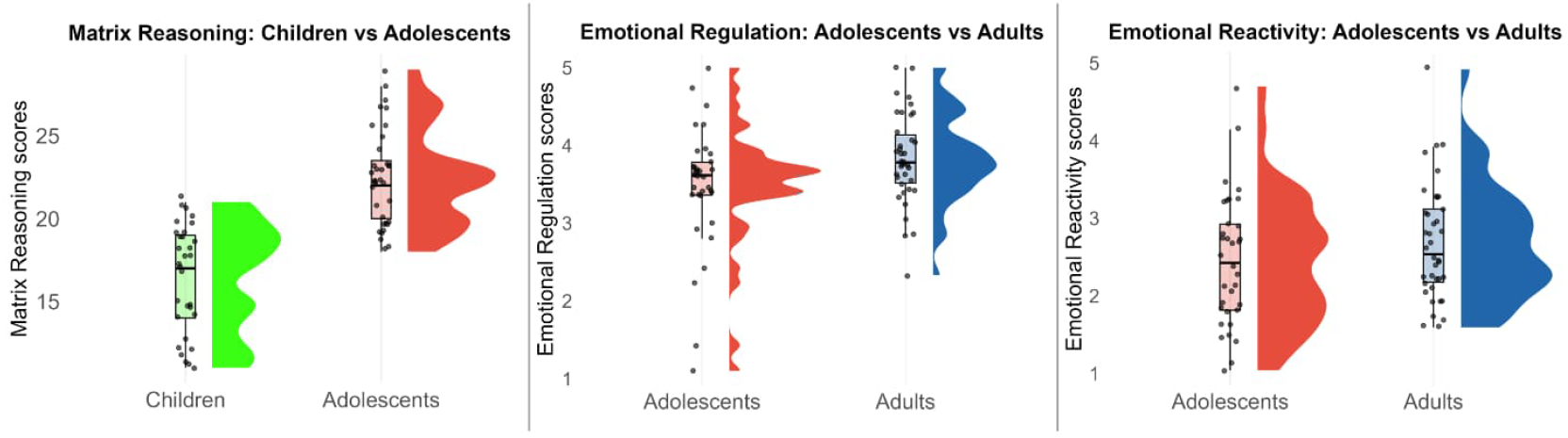
Cognitive and emotional scores across age groups. From left to right: Matrix Reasoning raw scores in children and adolescents, Emotional Regulation scores in adolescents and adults, Emotional Reactivity scores in adolescents and adults.

At the primary threshold (*p*_FWE_ < 0.05, 20-voxel cluster extent), no significant associations between cerebellum GMV and cognitive and emotional function were detected. Given the exploratory nature of these analyses, additional regression tests were run at *p* < .001 uncorrected (20-voxel extent). We report these results here with the caveat that findings should be considered preliminary until corroborated in additional studies. These exploratory analyses revealed both age group differences and within-group patterns linking cerebellar GMV to cognitive and emotional function (see Figure 8). For cognitive performance (Matrix Reasoning scores), adolescents displayed more positive associations with GMV in left Crus I and lobule VIIIa compared to children (8a, left panel). Within adolescents, there was an association between higher Matrix Reasoning scores and increased GMV in lobules VIIIa and VIIb (8a, right panel).

**Figure 8.**
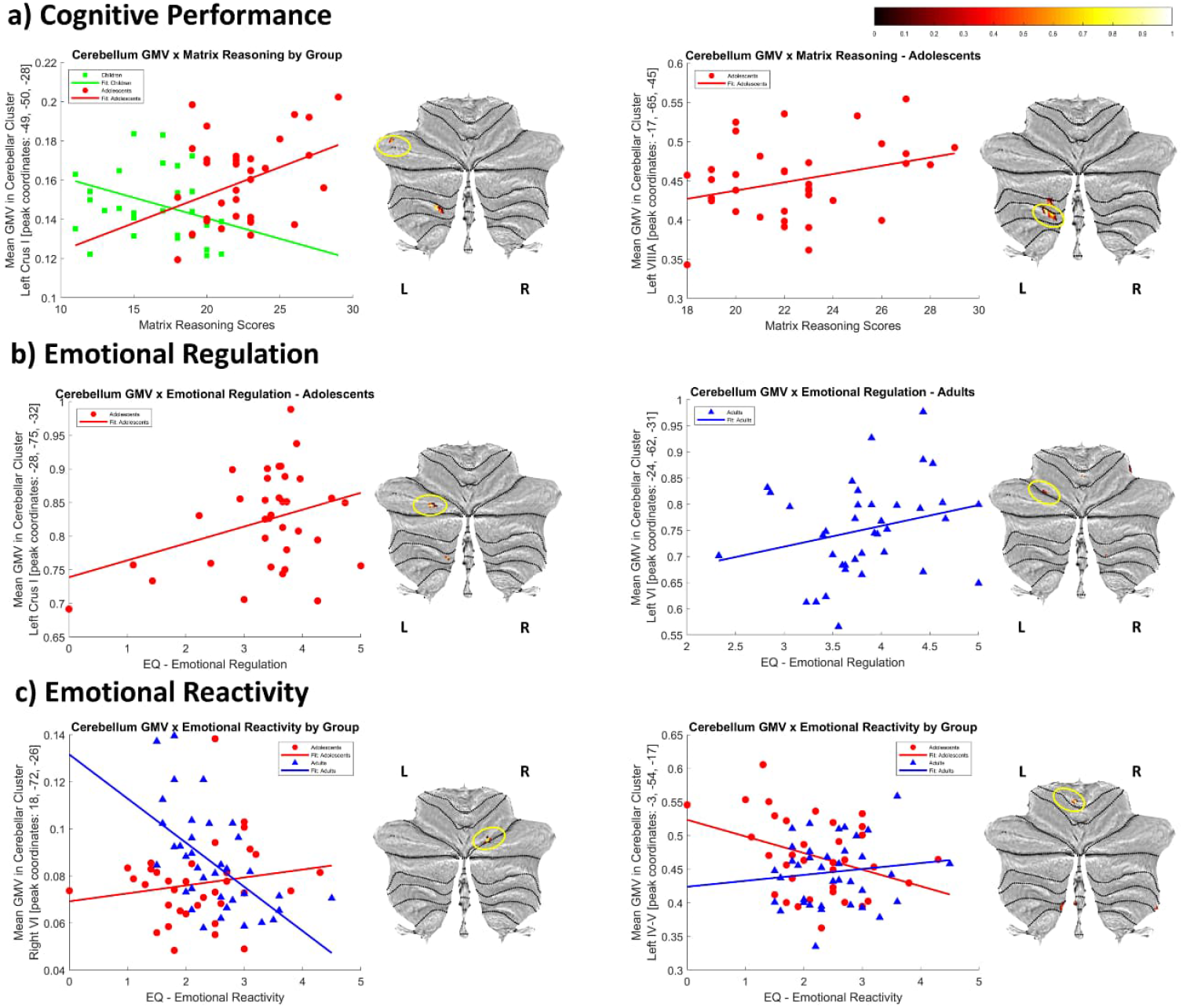
Exploratory associations between cerebellar gray matter volume (GMV) and cognitive and emotional function. Voxel-wise regression analyses at *p* < .001 uncorrected (20-voxel extent) showed potential associations linking GMV to **a) Cognitive Performance** (as indexed by Matrix Reasoning scores) in Crus I and lobule VIIIa, **b) Emotional Regulation** (Crus I, lobule VI), and **c) Emotional Reactivity** (lobules VI, IV-V). Plots show mean GMV (scaled 0-1, relative units from modulated images) for illustrated clusters indicated by yellow circles on SUIT flatmaps. Colorbar represents scaled *t*-values. Green squares and lines = Children; red dots and lines = Adolescents; blue triangles and lines = Adults. L = left; R = right.

For emotional regulation (EQ scores), we found positive associations with GMV in left Crus I and lobule VIIIa in adolescents (8b, left panel), and in lobules VI and VIIb in adults (8b, right panel), with no age-dependent differences. Finally, for emotional reactivity (EQ scores), adolescents compared to adults showed a positive association with GMV in right VI (8c, left panel), with an inverted pattern in left IV-V (8c, right panel). Within-group, emotional reactivity showed a positive association with GMV in left lobule VI in adolescents, and with GMV in left Crus I in adults (Supplementary Figures 4-5). Full cluster statistics are reported in Supplementary Table 1. For the sake of completion, we also report the results of age-invariant analyses at the *p* < .001 + 20 voxel extent threshold in Supplementary Figure 6. Briefly, age-independent negative associations were detected between cerebellum GMV in lobules VIIb and VIIIb for matrix reasoning scores and in Crus I and lobule V for emotional reactivity scores, while emotional regulation was positively related to GMV in lobule VIIIa and Crus I, and negatively related to GMV in Crus I and lobules V and VIIIb.

## 4. Discussion

Few studies have examined GMV of the cerebellum across development from childhood to adulthood, and to our knowledge, this is the first to apply cerebellum-optimized VBM to a sample spanning middle childhood (6 years) to mid-adulthood (40 years). In line with our hypothesis, we observed greater cerebellar GMV in adolescents compared to both children and adults, and in adults compared to children. This developmental pattern displayed regional specificity: some subregions showed early maturation with GMV stabilization from adolescence onward, while others followed an inverted U-shaped trajectory peaking in adolescence.

Our findings of inverted U-shaped patterns of voxel-wise GMV from childhood to adulthood in certain posterolateral regions (lobules VI and VIIb and Crus I) are partly consistent with previous studies performing cerebellar regional analyses (Romero et al., 2021; Tiemeier et al., 2010; Wierenga et al., 2014), albeit those relied on lower field strength MRI (1.5 T; Wierenga et al., 2014; Tiemeier et al., 2010) or mixed field strength datasets (Romero et al., 2021), which may limit spatial resolution. We extend previous findings with three important contributions. Firstly we show within-region specificity using VBM, allowing detection of age group differences that are not uniform or in entire cerebellar regions. Secondly, we identify posterolateral regions that appear to mature earlier (Crus I, lobule X), with stable GMV from adolescence to adulthood, and thirdly by showing vermal regions following as well an inverted U-shaped trajectory of GMV peaking in adolescence in vermis VI and VIIb. Importantly, this latter finding contrasts with Tiemeier et al. (2010) who reported no significant vermal GMV variations during adolescence, highlighting our study’s contribution to demonstrate adolescent-specific patterns of increased GMV in the vermis. Adolescent-related growth of these vermal regions are of particular interest given the affective nature attributed to the cerebellar vermis (reviewed in Adamaszek et al., 2017).

Adolescence is marked by acute neural remodeling, involving synaptic growth and pruning, myelination, and maturation of white matter tracts, all occurring under the influence of hormonal changes (e.g., Curtis et al., 2024), and accompanied by cognitive-emotional changes (e.g., Kolskår et al., 2018) which may both shape and be shaped by cerebellar regional development. In our study, adolescents had greater regional cerebellar GMV than both children and adults. Clusters of greater GMV spanned almost the entire cerebellar cortex compared to children, with the largest effects found in bilateral Crus II, right Crus I, right I-IV and bilateral lobule X. These findings align with Gaiser et al. (2024), who reported pronounced age-dependent volume growth from childhood to adolescence in posterior lobules. However, our results contrast with those of Bernard et al. (2015), who also reported inverted U-shaped developmental trajectories but with peak cerebellar volumes occurring around age 30. Extending previous literature, our results highlight adolescence as a sensitive window for cerebellar maturation.

The most pronounced GMV differences between adults and children were localized in the posterolateral cerebellum, specifically Crus I, Crus II, and lobules IX and X bilaterally. Out of those areas, tractography findings by Salmi et al. (2010) have linked Crus I and II to lateral prefrontal regions, with increased activation scaling with cognitive demand. Notably, adolescents, compared to adults, also showed a pattern of greater cerebellar GMV with peak coordinates in a cluster encompassing vermis VI, bilateral lobule VI, vermis VIIb and left VIIb. Functional MRI (fMRI) studies have previously linked these posterior cerebellar regions to emotional processing and learning (Kattoor et al., 2014; Pierce et al., 2023). These functions mature from childhood into adulthood and rely on coordinated activity between the cerebellum, prefrontal cortex, and emotion-related brain areas such as the amygdala (Gee et al., 2013). Adolescence is a particularly sensitive developmental window, marked by increased vulnerability to the onset of psychiatric disorders (Paus et al., 2008; Kessler et al., 2005). Importantly, structural and functional alterations in posterior cerebellar regions -such as those reported in the present study-have been consistently associated with affective (Depping et al., 2018; Xu et al., 2017; Shad et al., 2012) and anxiety disorders (Kolesar et al., 2019; Roy et al., 2013; Talati et al., 2013), highlighting the potential contribution of cerebellar maturation to the emergence of emotional dysregulation during this period.

Accordingly, our exploratory analyses pointed out both age-dependent and age-independent associations between cerebellar regional GMVs and cognitive and emotional function. Even though these findings did not survive correction for multiple comparisons, their overall pattern aligns with prior evidence of functional location of cognitive and affective processing found in the posterolateral cerebellum (Guell et al., 2018b). For cognition, adolescents displayed more positive associations than children between Matrix Reasoning scores and GMV in left Crus I and VIIIa, potentially suggesting increased recruitment of these regions during adolescence. For emotional functioning, associations involved left-lateralized regions (lobule VI and Crus I), which may tentatively align with recent evidence of hemispheric assymetries in cerebellar contributions for fear-related processing (Pakusch et al., 2025).

Additionally, patterns within adolescents indicated a potential functional heterogeneity with overlapping positive and negative associations with emotional function in Crus I. Emotional reactivity showed divergent associations: in adolescents, lobule VI was positively related to emotional reactivity, whereas in adults lobules IV-V, VIIIb, and Crus I were involved. Although lobules IV-V are classically of sensorimotor nature, prior fMRI evidence has linked them to negative affective processing (Park et al., 2010), potentially pointing to a dual role in adulthood. Negative age-invariant associations between emotional reactivity and GMV in lobule V and Crus I further suggest potential developmental functional shifts in cerebellar GMV recruitment. Taken together, these findings hint at dynamic, selective recruitment of cerebellar regional GMV across development, yet they should be interpreted with maximum caution given their exploratory, uncorrected nature.

### 4.1 Strengths and Limitations

While our study provides valuable insights into cerebellar morphological changes across childhood, adolescence, and adulthood, some limitations should be considered. First, the use of a cross-sectional design limits inferences about developmental trajectories; thus, replication in longitudinal cohorts would strengthen the interpretation of age-related patterns. Second, we did not collect handedness data, which might have aided interpretation of regional GMV. However, a recent study found no relationship between handedness and cerebellar GMV in adults (Kavaklioglu et al., 2017), and the implications of handedness for gray matter asymmetries remain inconclusive (Ocklenburg et al., 2016; Wang et al., 2021). Third, although the exploratory associations between cerebellar regional GMV and cognitive and emotional measures were discussed in line with previous studies, our findings were uncorrected and highlight the need for replication in larger, longitudinal samples.

Despite these limitations, several strengths should be emphasized. First, we applied a fully automated, cerebellum-optimized volumetric pipeline (ENIGMA Cerebellum Volumetrics), enhancing regional specificity and precision beyond what has been possible in other studies relying on whole-brain templates or predefined cerebellar regions. By spanning a wide age range (6–40 years), our design allowed for the detection of both region-specific nonlinearities and periods of stabilization in cerebellar GMV. We also conducted an extensive and detailed pre and postprocessing involving multiple levels of quality control, to ensure rigour and validity of the results. Furthermore, although the ACAPULCO algorithm was trained and developed with adult MRI data (Kerestes et al., 2022), our analysis of T1-w data from children and adolescents did not yield aberrant parcellations. This supports the pipeline’s performance, as the cerebellum typically reaches a structurally stable pattern during infancy (reviewed in Leto et al., 2016), with major morphological changes and rearrangements occurring in earlier stages than the mean age of the children group herein analyzed (8 years). Additionally, the methods and results adhered to established VBM reporting guidelines and good practices (Chen et al., 2017; Nichols et al., 2017; Ridgway et al., 2008), and facilitates transparency and reproducibility by employing an open-source neuroimaging pipeline.

### 4.2 Conclusion

In conclusion, this study reveals distinct patterns of cerebellar GMV across development, from middle childhood to adulthood, with certain posterolateral regions reaching mature GMV in adolescence, while others follow an inverted U-shape pattern peaking during this period. Notably, we extend previous work by adding within-region specificity and a larger age range, as well as showing transient GMV increases within cerebellar vermal regions during adolescence. Future studies should expand on these findings by investigating I) selective recruitment of cerebellar regions supporting cognitive and emotional functioning, and II) cerebellar-subcortical and cerebellar-cortical interactions across development to better characterize the functional role of the cerebellum and its connections in neurotypical development, as well as neurodevelopmental and adolescent-onset disorders.

## CRediT authorship contribution statement

**Richard Apps:** Writing – review & editing. **Andreas Frick**:, Supervision, Resources, Project administration, Investigation, Funding acquisition, Data curation, Conceptualization. **Matilda A. Frick**: Writing – review & editing, Investigation. **David Fällmar**: Writing – review & editing, Investigation. **Malin Gingnell**: Writing – review & editing, Resources, Project administration, Investigation, Funding acquisition, Conceptualization. **Patricia Gil-Paterna**: Writing – original draft, Visualization, Formal analysis, Methodology, Conceptualization, Data curation. **Johanna M. Hoppe**: Supervision, Writing – review & editing, Investigation, Conceptualization. **Dagmar Timmann:** Writing – review and editing, Conceptualization. **Ebba Widegren**: Writing – review and editing, Project administration, Investigation.

## Acknowledgements

The study was funded by a grant from Riksbankens Jubileumsfond to Dr. Frick and Dr. Gingnell (P17-0256:1). Dr. Frick was further supported by Kjell och Märta Beijers stiftelse and the Swedish Research Council (2021-03106). Dr. Fällmar was supported by the Swedish Society for Medical Research (SSMF, PD21-0136) and Hjärnfonden (PS2021-0026). This project received funding from the European Union’s Horizon 2020 research and innovation programme under the Marie Skłodowska-Curie grant agreement no. 956414. The computations and data handling were enabled by resources provided by the National Academic Infrastructure for Supercomputing in Sweden (NAISS), partially funded by the Swedish Research Council through grant agreement no. 2022-06725. The funding sources had no role in study design, data collection, analyses, publication decisions, or manuscript editing.

We thank Drs. Rebecca Kerestes and Srinivas Balachander for troubleshooting support with the ACAPULCO pipeline, Hampus Berg for scripting assistance, and Prof. Karin Brocki for providing study resources.

## Code and Data Accessibility

The customized scripts of the cerebellum pipeline used herein with the ACAPULCO algorithm, as well as the setup of the SPM12 batch parameters, may be accessible upon reasonable request. Due to the sensitive nature of the data collected in this study, participants were assured raw data would remain confidential and would not be shared. The raw data that has been used is thus confidential and will not be shared. Processed data will be shared upon reasonable request.

