## Supplementary Material for "Cerebellar Gray Matter Volume Changes Across Development: Posterolateral and Vermal Transient Increases during Adolescence"

**Patricia Gil-Paterna**

Uppsala University, Department of Psychology,

752 37 Uppsala, Sweden.

**Conflict of interest**

The authors declare no conflict of interests.

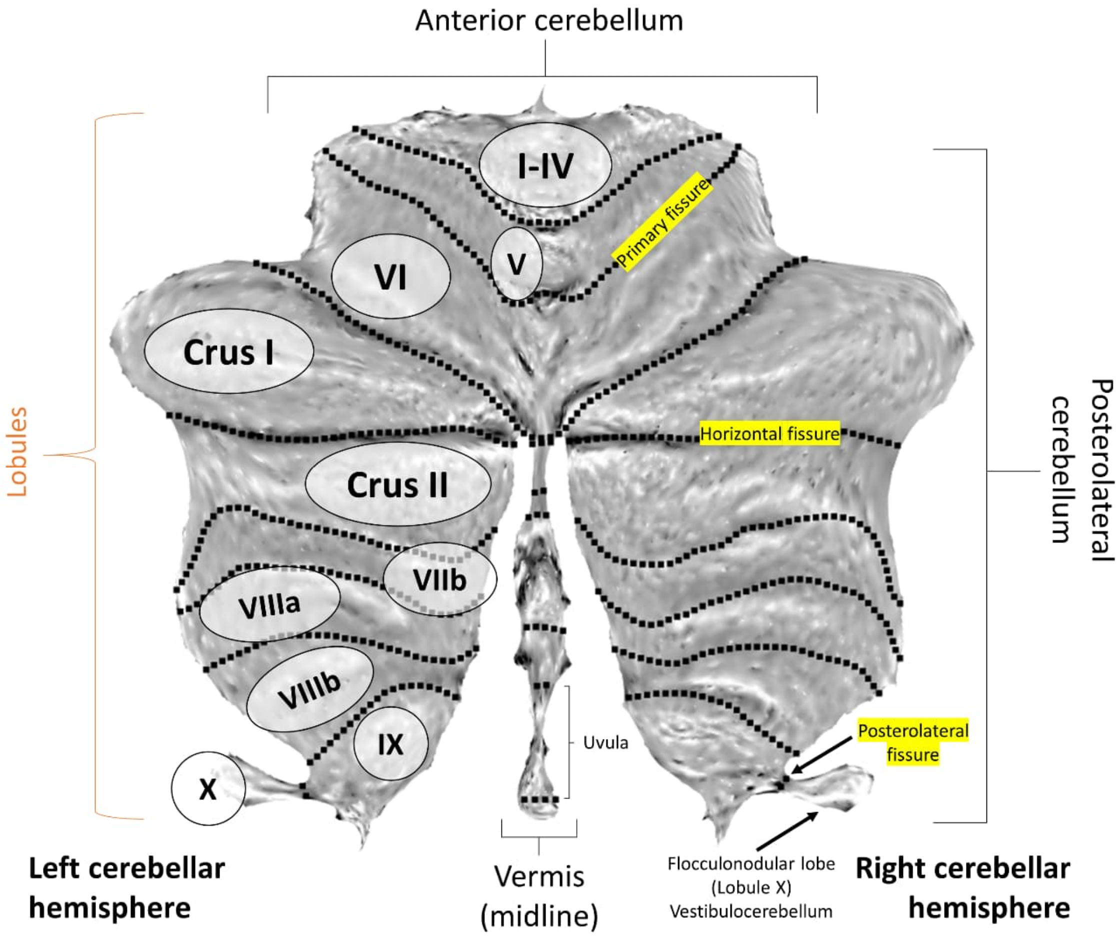


**Supplementary Figure 1. Anatomy and subdivisions of the human cerebellum.** The cerebellar cortex is shown with a schematic overall division: anterior lobe, posterolateral region, and midline vermis, from superior to inferior orientation. Lobules are labeled from I-IV to X in superior-to-inferior order. Structural boundaries between anterior and posterior cerebellum converge on lobule VI, which -though traditionally considered part of the anterior lobe- is increasingly included as part of the posterior cerebellum. Note that Crus I and Crus II together form the lobule VIIa, which is then followed by VIIb. Lobules labeled in the left cerebellar hemisphere are homologous to those in the right. The cerebellum flatmap was generated using the SUIT toolbox for SPM12 (from Diedrichsen & Zotow, 2015).

### Supplementary results

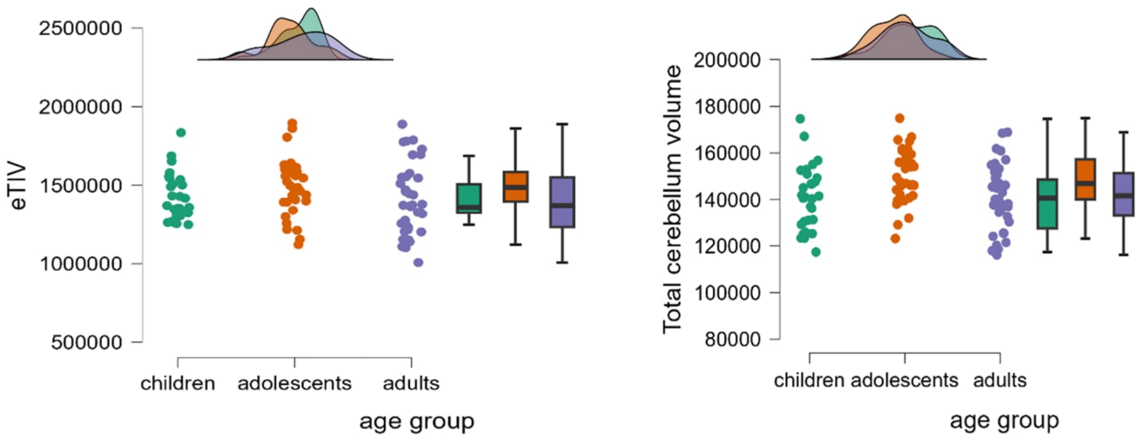


**Supplementary Figure 2. Brain volumetrics across age groups.** Boxplots displaying volumetrics for estimated total intracranial volume (eTIV) (left) and total cerebellum volume (right) across children, adolescents, and adults. Both eTIV and total cerebellum volume are expressed in $mm^3$.

Mean eTIV was 1420 ± 143.83 (SD) $cm^3$ in children, 1487 ± 179.17$cm^3$ in adolescents, and 1402 ± 223.92 $cm^3$ in adults. ANCOVA post hoc tests revealed that adolescents had significantly greater eTIV compared to adults (Cohen's $d$=.65; $p_{tukey}$= .017), Corresponding mean total cerebellum volumes per group were 139.91 ± 13.78 $cm^3$, 149.13 ± 11.61 $cm^3$, and 141.44 ± 14.02 $cm^3$, respectively. Thus, total cerebellum volume as a percentage of eTIV was 10.15% in children, 9.97% in adolescents, and 9.92% in adults. Children had significantly smaller cerebellum eTIV-corrected volume compared to adolescents (Cohen's $d$=-.61; $p_{tukey}$= .04). Data were approximately normally distributed with minor outliers over the extreme quantiles. Levene's test showed that homogeneity of variances was met for both eTIV and total cerebellum volume across groups. Within-group analyses showed no significant sex differences in eTIV or total cerebellum volume.

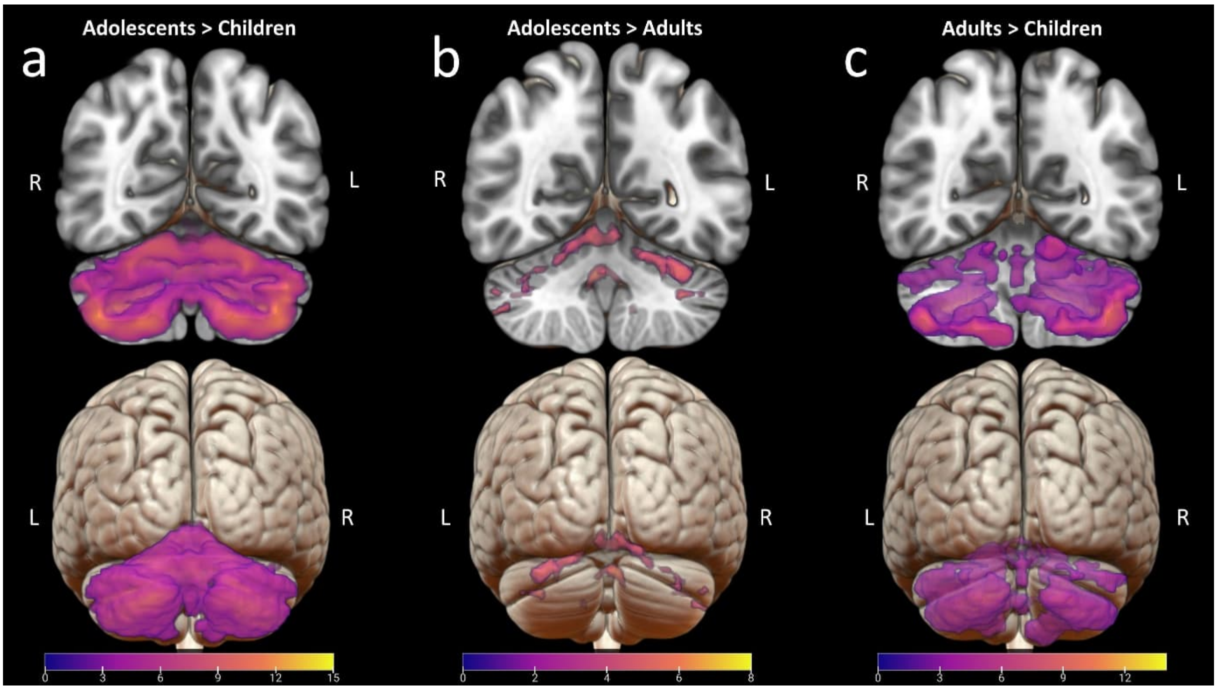


**Supplementary Figure 3. Voxel-based morphometry differences between age groups in gray matter volume. a) Adolescents > Children:** cerebellar regions with significantly greater gray matter volume (GMV) in adolescents compared to children. **b) Adolescents > Adults**: cerebellar regions with significantly greater GMV in adolescents compared to adults. **c) Adults > Children**: cerebellar regions with significantly greater GMV in adults compared to children**.** Statistical T-maps thresholded at $p_{FWE}$<.05 and cluster extent 20 voxels were overlaid on MNI152 brain template using MRIcroGL (Chris Rorden; https://github.com/rordenlab/MRIcroGL). Colorbars represent *t*-values. Anterior orientation is displayed above, while posterior is shown below. L = left; R = right.

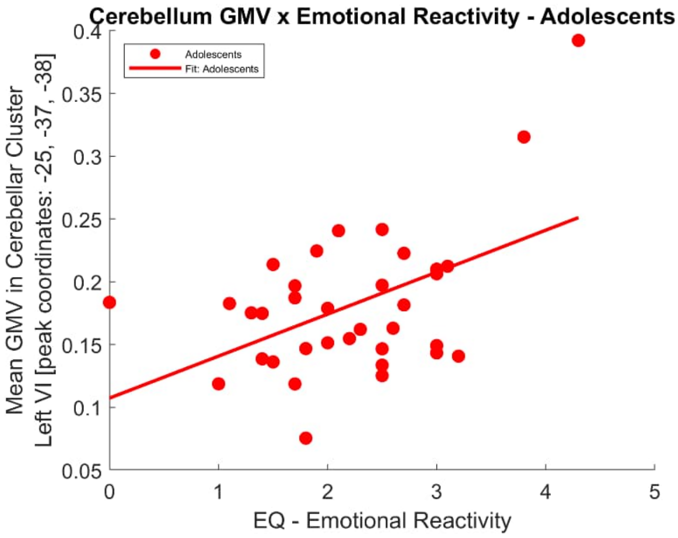


**Supplementary Figure 4.** Regression plot illustrating the relationship between gray matter volume (GMV) in a cluster within the left lobule VI and emotional reactivity scores from the Emotion Questionnaire (EQ), in adolescents.

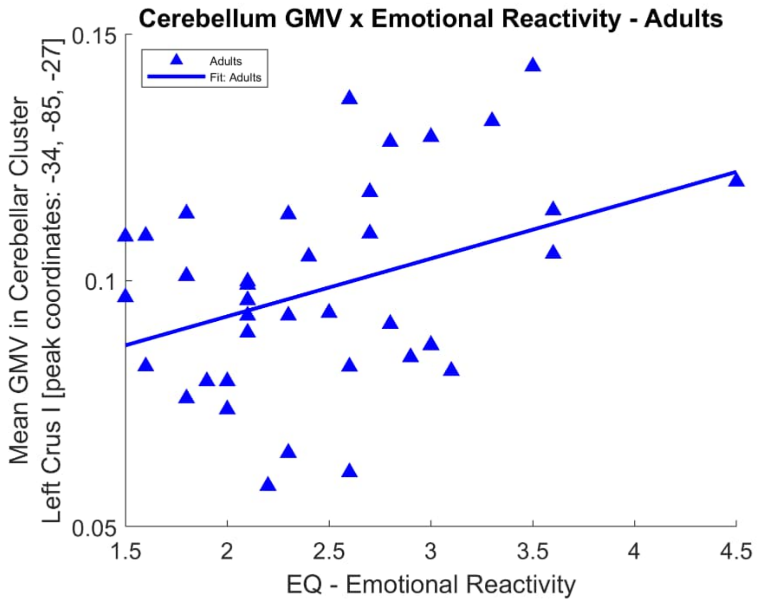


**Supplementary Figure 5.** Regression plot illustrating the relationship between gray matter volume (GMV) in a cluster within the left Crus I and emotional reactivity scores from the Emotion Questionnaire (EQ), in adults.

**Supplementary Table 1. Results from voxel-based morphometry of associations between cerebellar gray matter volume (GMV) and cognitive and emotional functioning.** Thresholds set to uncorrected p<.001 and cluster extent of 20 voxels (20 mm$^3$).

| **Cerebellar region** | **Peak MNI coordinates x y z** | **Volume (mm³)** | ***t*** | ***p (uncorrected)*** | **Direction + positive - negative** |
|---|---|---|---|---|---|
| **Cognitive Performance** | | | | | |
| ***Adolescents > Children*** | | | | | |
| Left Crus I | -49 -50 -28 | 22 | 3.88 | < *.001* | + |
| Left VIIIa | -16 -65 -46 | 24 | 3.75 | < *.001* | + |
| ***Adolescents*** | | | | | |
| Left VIIIa | -17 -65 -45 | 103 | 4.42 | < *.001* | + |
| Left VIIb | -16 -71-43 | 39 | 4.30 | < *.001* | + |
| **Emotional Regulation** | | | | | |
| ***Adolescents*** | | | | | |
| Left Crus I | -28 -75 -32 | 32 | 3.95 | <.001 | + |

| | | | | | |
|---|---|---|---|---|---|
| Left VIIIa | -9 -70 -53 | 20 | 3.75 | <.001 | + |
| ***Adults*** | | | | | |
| Left VI | -24 -62 -31 | 23 | 3.67 | <.001 | + |
| Right VI | 35 -41 -33 | 21 | 3.33 | <.001 | + |
| Right VIIb | 14 -74 -57 | 24 | 3.89 | <.001 | + |
| **Emotional Reactivity** | | | | | |
| ***Adolescents > Adults*** | | | | | |
| Right VI | 18 -72 -26 | 47 | 3.46 | <.001 | + |
| Left IV-V | -3 -54 -17 | 34 | 4.37 | <.001 | - |
| ***Adolescents*** | | | | | |
| Left VI | -25 -37 -38 | 28 | 4.23 | <.001 | + |
| ***Adults*** | | | | | |
| Left Crus I | -34 -85 -27 | 22 | 3.99 | <.001 | + |

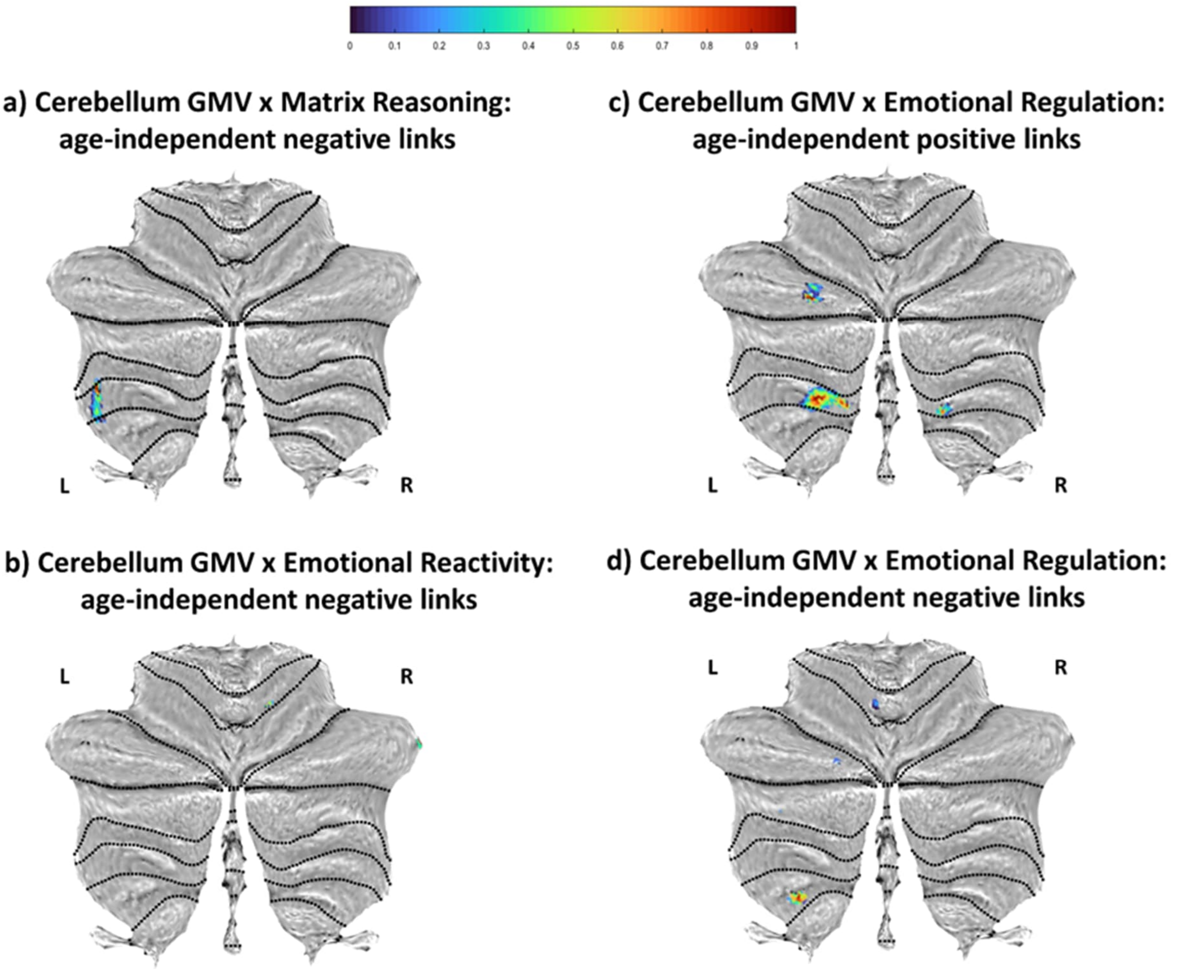


**Supplementary Figure 6. Age-independent associations between cerebellar gray matter volume (GMV) and cognitive and emotional function.** Age-independent negative associations were detected at the p<.001 uncorrected level with cluster extents ≥20 voxels between cerebellum GMV in **a**) lobules VIIb and VIIIb for matrix reasoning scores and. **b**) in Crus I and lobule V for emotional reactivity scores. **c**) Emotional regulation was positively related to GMV in lobule VIIIa and Crus I, and **d**) negatively related to GMV in Crus I and lobules V and VIIIb. Colorbar (top center) represents scaled t-values (0-1) from *t*-maps thresholded at *p* <.001 (uncorrected) with extent threshold of 20 voxels. Clusters are overlaid on cerebellum flatmaps. L = left; R = right.
